# 3D-printing fabrication of microwave-microfluidic device for droplets network formation and characterisation

**DOI:** 10.1101/2024.04.08.588546

**Authors:** Kai Silver, Jin Li, Pantelitsa Dimitriou, Colin Kallnik, Adrian Porch, David Barrow

## Abstract

Microwave-microfluidic devices (MMDs) have emerged as precision tools for the rapid, accurate, sensitive, and non-invasive characterisation of low-volume liquids. However, the fabrication of MMDs remains a significant challenge due to the complexities associated with integrating fluidic ducts and electronic components. Herein, we present a versatile and economical 3D-printing approach for MMD fabrication, using liquid metal as an electrical conductor. Cyclic olefin copolymer, polylactic acid and polypropylene were identified as potential printable dielectric materials for MMD fabrication. 3D-printed cyclic olefin copolymer substrates exhibited the lowest loss tangent of 0.002 at 2.7GHz, making it an ideal material for high frequency engineering. Liquid metal, specifically gallium indium eutectic, was injected into the printed ducts to form conductive microwave structures. Exemplar MMDs were fabricated to integrate split-ring type microwave resonators and droplet-forming fluidic junctions. These devices were applied in the formation and characterisation of water-in-oil emulsions for constructing definable lipid-segregated droplet interface bilayer (DIB) networks. This work not only indicates the feasibility of using 3D-printing for rapid prototyping of customised MMDs but also demonstrates the potential of MMDs as a new research tool for biochemistry and synthetic biology.

## Introduction

Microwave-microfluidic devices (MMDs) represent a class of integrated instruments for processing and analysing liquid samples and soft matter ^1–4^. Droplet microfluidics has significantly advanced in the last three decades, bringing capabilities of precision liquid handling and high-throughput experimentation using droplet-based materials ^5,6^. Meanwhile, microwave sensors have emerged as a technology for the rapid and sensitive characterisation of materials, primarily by measuring their dielectric properties^7,8^. The synergistic integration of microwave and microfluidic technologies enables highly accurate, non-invasive measurements of emulsions ^9^. MMDs also provide rapid dielectric heating of biological samples in an aqueous environment, including DNA plasmids and living cells ^10–12^. Recent developments in MMDs have extended their application to biosensing, notably in monitoring blood glucose levels ^13^, underpinning the potential of MMDs for innovations in analytical chemistry, biotechnology, pharmaceuticals, and medical diagnostics.

The fabrication of MMDs can be technologically complex due to the intricate micromachining of fluidic and microwave structures within the same microsystem^14^. The design of microfluidic and electronic components, along with the selection of appropriate microwave substrate materials, is critical for MMD functionality and performance. The conductive component in MMDs creates electrical pathways for microwave transmission, while dielectric substrates shape and confine the electric field, directing microwave energy to the samples. The substrate materials need to be machinable to create micro-scale fluidic ducts for liquid flow. Typical substrates materials for constructing microfluidic devices include epoxy-glass laminates, polymers, silicon, glass, metals, and ceramics, due to their compatibility with microfluidics. However, in general, the integration of microwave functionality with microfluidic functionality, requires more stringent requirements of the constructional material (e.g. Teflon-based printed circuit boards^15^) to minimise dielectric loss.

Recently, a form of additive manufacturing known as 3D printing technology, has been applied to the fabrication of micro-scale devices^16^, offering rapid prototyping of complex integrated structures. This manufacturing approach enhances the flexibility of device design and enables quick turnaround from concept to prototype. Multiple materials, such as both dielectric and conductive filaments, can be 3D-printed on the same prototype, allowing for the fabrication of monolithic droplet-forming and sensing microfluidic devices with polymeric fluidic channels and integrated electrodes ^17,18^. Notably, liquid metals, such as gallium-indium eutectic, have been routinely used to form conductive parts in microfluidic devices ^19^. Liquid metal can be injected into void channels to form irregular shaped, spiral electrodes within 3D-printed microchip electrophoresis devices ^20^. 3D printing technologies have also been applied to microwave instrumentation, such as waveguides, resonators, and antennas ^21–23^, incorporating liquid metals as a tool to impose microwave properties on the devices ^24^. Additionally, planar microwave resonators have been attached to 3D-printed microfluidic devices for the analysis of flowing liquid materials ^25^. However, the use of liquid metals has not replaced conventional, costly, and complex manufacturing methods for the electrodes. Therefore, the potential of 3D printing for MMD fabrication remains largely unexplored to date, despite its promise for enhancing accessibility and customisability in various applications.

In this context, we introduce a novel and facile fused filament 3D-printing method for MMD fabrication, incorporating gallium-indium eutectic liquid metal. Various 3D-printable substrate materials and printing settings were evaluated to optimise MMD performance. A comprehensive, step-by-step protocol is provided for the design and fabrication of MMDs. The demonstrated functional structures inside MMDs include split-ring type microwave resonators, droplet-forming fluidic junctions, and droplet-handling fluidic circuits. The printed MMDs were used to generate and characterise single emulsions within flowing ducts. The microwave signatures correlated with the formed droplet sizes, which was consistent with corresponding optical measurements. Subsequently, the MMDs were used for the formation of lipid-segregated droplet interface bilayers (DIBs) networks, and the number and the size of droplet components within a network were identified from the microwave readout. DIB networks have wide applications in molecular biology ^26,27^, artificial cellular membranes ^28^, and the engineering of bottom-up synthetic cells and tissues ^29,30^. The presented 3D printing approach of MMDs are highly accessible and has the potential to broaden their applications in miniaturised systems for potential bioengineering, biochemical and soft material research.

## Experiments

### Materials

Ultimaker polylactic acid (PLA) and Ultimaker polypropylene (PP) filaments were purchased from CREATE Education Projects Ltd. Cyclic olefin copolymer (COC) filaments were purchased from CREAMELT. Plastic syringes (BD Plastipak) were purchased from Fisher Scientific. Microfluidic tubing (O.D. ᴓ = 1.58 mm, I.D. ᴓ = 0.50 mm, Teflon), polyether ether ketone (PEEK) microfluidic fittings and connectors were purchased from Cole-Parmer UK. Gallium-indium eutectic and all the chemicals were purchased from Merck. Phospholipid 1,2-diphytanoyl-sn-glycero-3-phosphocholine (DPhPC), hexadecane and silicone oil AR20 were purchased from Merck. A DPhPC-containing oil phase (60% hexadecane and 40% silicone oil AR20) was prepared at a concentration of 8 mg/mL, following an existing published protocol^31^. Sub-miniature version A (SMA) connectors, conductive copper films and soldering wires were purchased from RS Components UK. All the materials were used as delivered unless otherwise mentioned.

### Microwave-microfluidic device design and modelling

MMD structures were designed using the computer-aided design interface of COMSOL Multiphysics software. The geometry files built in COMSOL were exported as STL files. The electromagnetic field within the MMDs was simulated using the RF module of COMSOL Multiphysics software (version 5.6). MATLAB 2023b, in conjunction with the signal processing toolbox, was used to post-process numerical data.

### Microwave-microfluidic device fabrication

The geometry STL files of MMD were sliced to G-Code files using Ultimaker CURA software (version 4.3.0). All printing profiles were used as the materials default setting, unless otherwise specified, with the printing layer thickness set at 0.1 mm. The MMDs were fabricated using Ultimaker S5 Pro Bundle 3D printers, following the suppliers’ suggestions on printing setup. The ambient humidity was regulated by a domestic dehumidifier during 3D printing. The printed devices were stored alongside silica gel to avoid moisture inside a box before use. The assembly of MMDs is detailed in the results section of this paper.

### Microwave characterisation

The microwave permittivity of 3D-printed plastic sheets was measured using a split-post dielectric resonator, operating at 2.7 GHz, and an Agilent E5071B Vector Network Analyser (VNA). S_21_ measurements were made to determine the resonant frequency and quality (Q) factor with and without a sample. Cavity perturbation theory was then applied to extract the complex permittivity ε, where the real part (ε _1_) being the polarisation term and the imaginary part (ε _2_) the loss term. The samples were printed as 50x30x0.5 mm sheets. Both line and triangular printing formats were explored, with the infill percentage set to 100% (later adjusted to 50% for COC). Both line and triangular printing formats were explored. A Mitutoyo micrometer was used to measure all sheet thicknesses.

The permittivity and S_21_ parameters of various solvents were measured using a cylindrical cavity resonator, operating in the TM_010_ mode at 2.4GHz ^32^, and the VNA. Liquids were held in a 2.5mm inner diameter, 3mm outer diameter clear FEP tube (FEP has a very low loss tangent which was negligible in these experiments). The tube was placed along the axis of the cavity, where the microwave electric field was highest and parallel to the tube. The obtained values were used to design the microwave structures of the MMDs.

### Microfluidic setup

Positive displacement syringe pumps (KD Scientific Legato 210) were used to inject fluids into the MMDs, via syringes, connectors, and tubing. The connections of tubing to MMDs were sealed with UV adhesive (Permabond) using a 365nm UV torch. A picture of the microwave-microfluidic experimental setup is shown in Fig S1. The input flow rate of fluids was controlled by the syringe pumps with actuation governed by customised Python code to program droplet formation. Dispersed aqueous phases were injected into either T-shaped or flow-focusing droplet-forming junctions, broken into discrete droplets by a mineral oil continuous phase, and then flowed through the gap region of the split-ring microwave resonator designed within the 3D-printed MMDs.

### Imaging, data collection and analysis

All the experimental photos and videos were captured using a USB microscope (Dino-Lite, AM7915MZTL) or a Nikon MM800 optical measurement stereomicroscope. An E5071B VNA was used to obtain the microwave signature, controlled using a NI GPIB-USB-HS card and Python code utilising the PyVISA package. The size measurement of microfluidically-formed droplet was analysed using ImageJ software. The data were processed with Microsoft Excel software before being analysed using MATLAB 2021b with the signal processing toolbox 8.7 and curve fitting toolbox 3.6.

## Results

### Microwave-microfluidic device design

**Error! Reference source not found**. presents an example of a 3D-printed microwave-microfluidic device. The device structure includes a fluidic manifold with various channels for the purposes of the fluid conduction and the incorporation of microwave feeds. The substrate material was a 3D-printed dielectric plastic block. The ring resonator, a square-faced torus made of EGaIn liquid metal, was designed to achieve a fundamental resonant frequency of 2 GHz. This ring resonator had an inner radius of 7 mm, an outer radius of 8 mm, a height of 1 mm, and a gap width of 1.4 mm. Electromagnetic energy entering and exiting the resonant structure was coupled using two SMA connectors. The end of the SMA centre pin produced an electric field that coupled into the ring. COMSOL Multiphysics was used to optimise the design of the ring resonator for efficient microwave sensing, transferring and storing energy from the VNA into the resonator, to interact with the sample in the gap region. The VNA continuously measured the voltage reflection coefficient magnitude, S21, in the frequency domain. The resonator’s performance was informed by extracting the resonant frequency (f0), quality factor (Q), and insertion loss (L the peak of S21 at resonance) from the VNA’s readings.

The fluidic manifold included fluidic inlets, droplet-forming fluidic junctions, circuits, and outlets. An expansion duct directed the formed droplets and continuous carrier phase, to flow through the gap region of the split-ring structure to enable the microwave sensing. At around 2 GHz resonant frequency, water retains a very high relative permittivity of 80 ^33^, whereas mineral oil phase and air have much lower permittivities of ε_mineral_=3.29 and ε_air_=1, respectively. Therefore, the water phase has a high induced electric dipole moment compared to mineral oil and air. As a water droplet enters the gap region, more electrical energy is stored within the gap, causing f_0_ to decrease. Additionally, due to dipole relaxation of water molecules, the quality factor (Q) decreases. Monitoring these changes provides real-time sensing of dispersed water droplets in continuous low-permittivity oil phases. An example of microwave signal detection is shown in Figure 1C. When the gap region was filled with low-permittivity mineral oil, the S21 measurement of the resonance frequency stayed at ∼ 2.2108 GHz. As a liquid droplet entered the gap region, the effective permittivity within the gap increased, leading to a decrease in resonant frequency to 2.2095 GHz, due to the greater stored energy. Such microwave signatures of water droplets flowing in the oil phase are volumetric measurements and provide an approach to characterise the liquid-based matrices and their morphology according to their electrical properties and flow profiles within the enclosed micro-ducts, using the MMDs.

**Figure 1.**
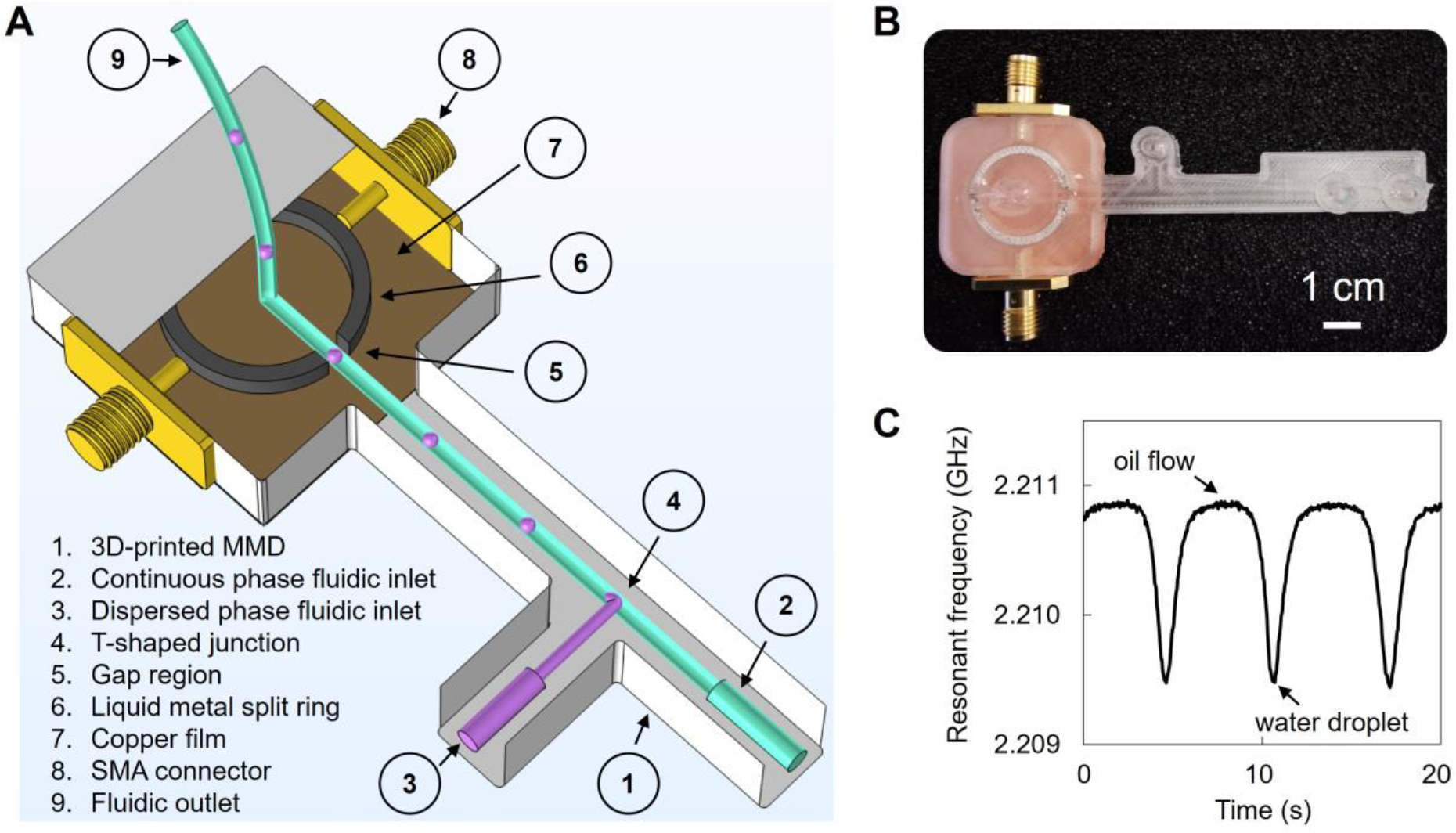
3D-printed microwave-microfluidic device (MMD) for droplet forming and detection. **>A**. Schematics of a 3D-printed MMD with labelled features. In this illustration, dispersed phase droplets are formed at a T-shaped droplet forming fluidic junction within a continuous phase fluid flow. The droplets pass though the gap region of a split-ring microwave resonator, created by liquid metal injection into 3D-printed ducts. The microwave resonator is activated by external microwave excitation through attached SMA connectors and grounded by a copper film which was adhered to the underside of the MMD. **B**. A sample of an assembled 3D-printed MMD. **C**. An example of a microwave signature when water droplets flow through the gap region of the split-ring microwave resonator. The water and oil phases exhibit different dielectric properties, leading to different microwave resonant frequencies, represented as peaks and troughs in the trace.

### 3D-printed MMD fabrication and optimisation

A step-by-step guide for manufacturing the 3D-printed MMD is presented in Figure 2. After fabricating the device substrate with a fused filament 3D printer, copper film tape was adhered to the bottom of the microwave structures to form a ground plane for the electric field. Two SMA connectors were inserted into the sides of the microwave structure. The PTFE dielectric shielding of the SMA connector pin was removed, allowing the pin to be positioned 1 mm away from the ring resonator. Adjusting the pin’s proximity to the resonator, without contacting the conductive part, alters the microwave energy coupling between the external source and the ring resonator. The SMA connectors, positioned 1 mm from the ring, resulted in a transmission loss of -9.1 dB at resonance. These connectors were soldered to the copper film. Liquid metal gallium indium eutectic was then injected into the microwave structures through open ports, forming the conductive paths. The open ports, fluidic inlets, and outlet connections were sealed with UV-curable adhesive to prevent leakage of the liquid metal and fluids. After assembly, the MMD was connected to a VNA machine for microwave signal transmission and sample measurements. For the droplet detection, fluids were injected to the MMD using syringes, via tubing and connectors by syringe pumps. As the droplets passed through the microwave structure, they were detected by the VNA, and imaged using an optical microscope.

**Figure 2.**
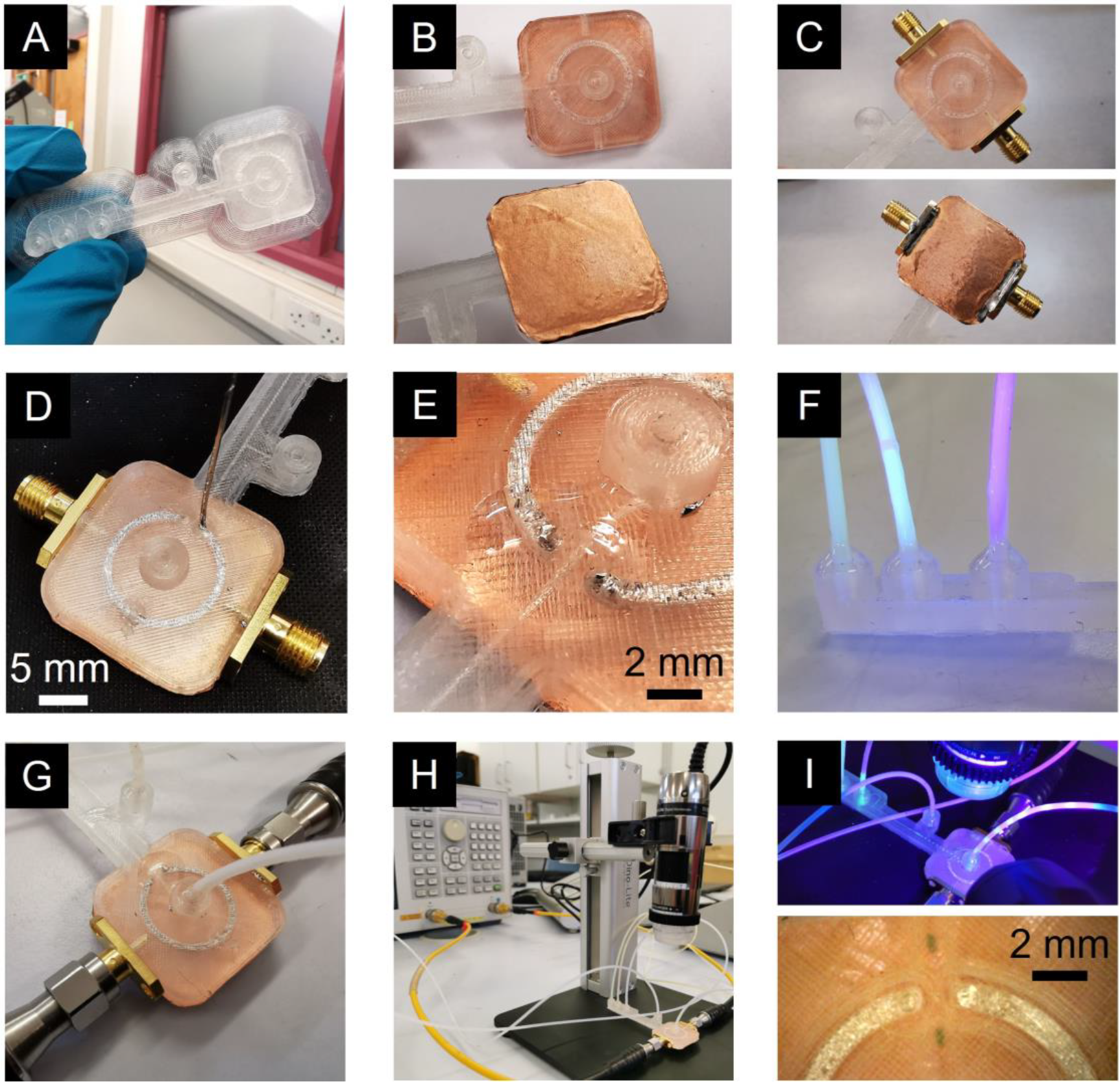
Step-by-step guide for manufacturing, assembly, and operation of 3D-printed MMD. **>A**. The chassis of MMDs were fabricated using fused filament 3D printing. **B**. A conductive ground plate (brownish, copper film) was then added below the planned position of the ring structure to reduce radiation losses and maintain a high Q-factor. **C**. Two SMA connectors were fixed in place and soldered to the ground plate. **D**. Liquid metal, gallium indium eutectic (EGaIn), was injected to the void channel of the MMD chassis through open ports using a glass syringe and metal needle, forming the conductive split-ring structure as the microwave resonator. **E**. The open ports were sealed with UV adhesive and cured with a UV torch to prevent leakage of EGaIn and preserve the integrity of the microwave structure features. **F**. The assembled MMD was connected to fluidic tubing (also sealed by UV adhesive) and syringe pumps. **G**. Electrical cables were attached via the SMA connectors to a network analyser. **H**. A USB microscope was utilised to observe the droplet formation. A VNA was used to send microwave signals and monitor the dispersed aqueous droplets and segment (blue) as they flowed in oil phase through the EM field at the gap region of the resonator inside the MMD, shown in **I-top** and **I-bottom**.

Injecting liquid metal into duct structures enables the patterning and encapsulating of reconfigurable electrical elements, such as electrodes^34^, sensors^35^, antennas^36^, and interconnects ^37^. In our fabrication approach for the MMD, gallium-indium eutectic (EGaIn) served as the conductive material, essential for transmitting microwaves. EGaIn’s electrical conductivity is 3.6 × 10^6^ S/m at room temperature, which is only one order of magnitude lower than that of noble metals^38^, and sufficient for microwave transmission. EGaIn is notably more dense, more cohesive and thermally conductive than water^19^. Also, at room temperature, its viscosity is only slightly higher, making it suitable for injection into circular micro-scale channels to form functional microwave components, within the 3D-printed devices.

The constructional substate materials of the MMDs must be of a suitably low microwave loss for appropriate microwave component performance. To assess the dielectric properties of 3D-printed substrates, plastic sheets were printed and measured to determine their complex permittivity ε, using a split-post dielectric resonator ^39^, as shown in Fig S2. This complex permittivity comprises a real part (ε_1_) for polarisation, and an imaginary part (ε_2_) indicating microwave loss. The polarisation term reflects the charge polarisation under an external electric field, while ε_2_represents losses in dielectric materials due to dielectric and conduction losses. Precise measurements were facilitated by aligning the electric field in the resonator’s gap region parallel to the plastic sheet surfaces. The real and imaginary permittivity components were deduced using cavity perturbation equations 1 and 2, where *G* = 2.20 ± 0.01 m^−1^ is the resonator constant, with the resonator constant being derived from EM field simulations or by measuring a known sample, such as PTFE. In equations 1 and 2, *t* is the plastic sheet thickness, and the subscripts “s” and “0” are measurements of resonant frequency *f* and quality factor *Q*, with and without sample, respectively.

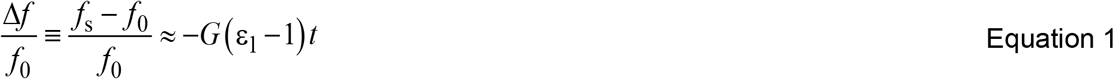

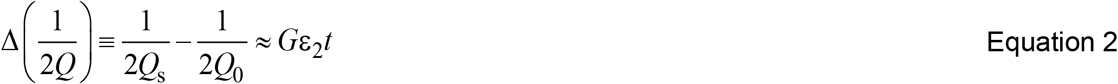

Table S2 shows the complex permittivity of the various printed plastic sheets using different fabrication conditions. Measurement accuracy is primarily affected by the systematic error in sheet thickness (± 50 μm) and its uniformity. COC demonstrated superior dielectric properties amongst the tested materials, exhibiting low polarisation, and minimal loss which is crucially for high Q. Tests at different infills showed that lower infill COC is preferable for high-frequency systems in certain applications ^40,41^, despite the potential for increased leakage of microfluidic flows, due to the air-plastic interfaces.

COMSOL simulations revealed the electric field at the gap region of the split-ring, as displayed in Figure 3A. The ring’s domain conductivity was set to that of EGaIn and the substrate to 100 % infill COC with the complex permittivity taken from Fig S2. The simulation results were consistent with the experimental microwave measurements. A broad sweep with a 1 GHz span about the SRR resonance was used to generate S-parameters for the 2-port SRR microwave device. The simulated S_21_ data were then imported into a MATLAB program, where Lorentzian curve fitting was used to extract the resonant frequency and *Q*. The shape of microfluidic chambers and ducts through the split-ring gaps influenced the electric field. Under simulation, sharp corners of the ducts can enhance the peak electric field compared to rounded corners (Figure 3Aiii maybe?). The precision fabrication of fluidic structures was limited by the printing resolution of the 3D printers employed.

**Figure 3.**
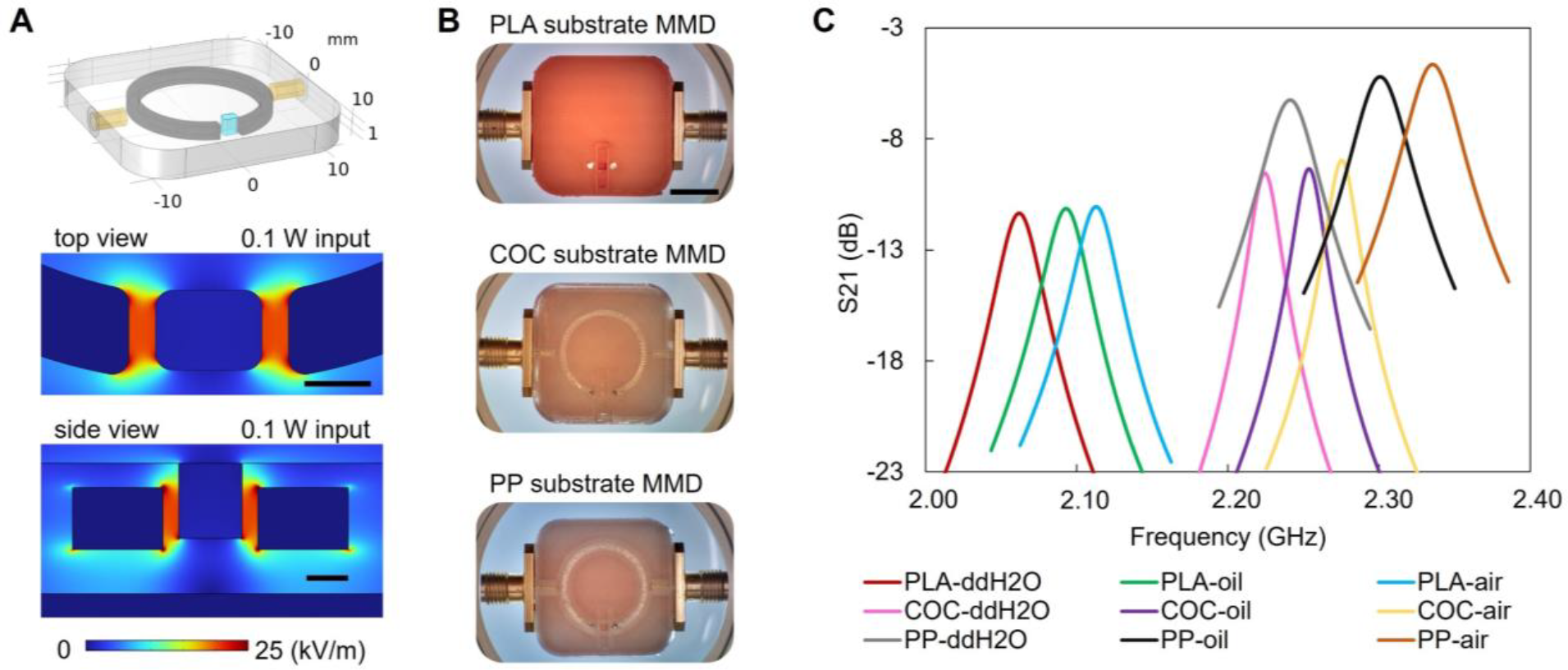
Performance of MMDs with different 3D-printed dielectric substrate materials in the microwave characterisation of liquid samples. **A**. 3D finite element modelling of the electric field at the gap region of split-ring microwave resonator using COMSOL Multiphysics. Scale bars denote 1 mm. Top-3D design of the split-ring; Middle and bottom-E-field distribution contours of the top view and sideview of the split-ring gap. **B**. Samples of MMDs made from dielectric PLA (top), COC (middle) and PP (bottom) filaments. All MMDs were built from the identical geometrical design and printing settings, with a layer thickness of 0.1 mm and line width of 0.23 mm. ===== Among these, the COC MMD exhibits superior microwave features, while PLA and PP are chosen for their chemical and physical properties suitable to various microfluidic applications. Scale bar denotes 1 cm. **C**. Microwave signatures obtained from using different MMDs to characterize various fluid materials within the gap region of the split-ring resonator.

As shown in Figure 4 A, MMDs with identical designs were fabricated using different dielectric substrates. COC and PP offered optical transparency, chemical resistance, and low permeability of gas and liquid, making them suitable for various biomedical and microfluidic applications^42^. The 3D-printed PLA had a higher loss tangent but remained partially viable for microwave sensing. PLA is a common 3D-printing material and widely adopted in many 3D-printing applications ^43^. Nylon printings were hydrophilic and has numerous applications in devices for life sciences and biomedicine, such as cell strainers, tubes, and dental dentures^44^. However, 3D-printed nylon has a relatively high microwave loss rendering it unsuitable for MMDs, because of the resultant low Q factor and sensitivity of a constructed device.

**Figure 4.**
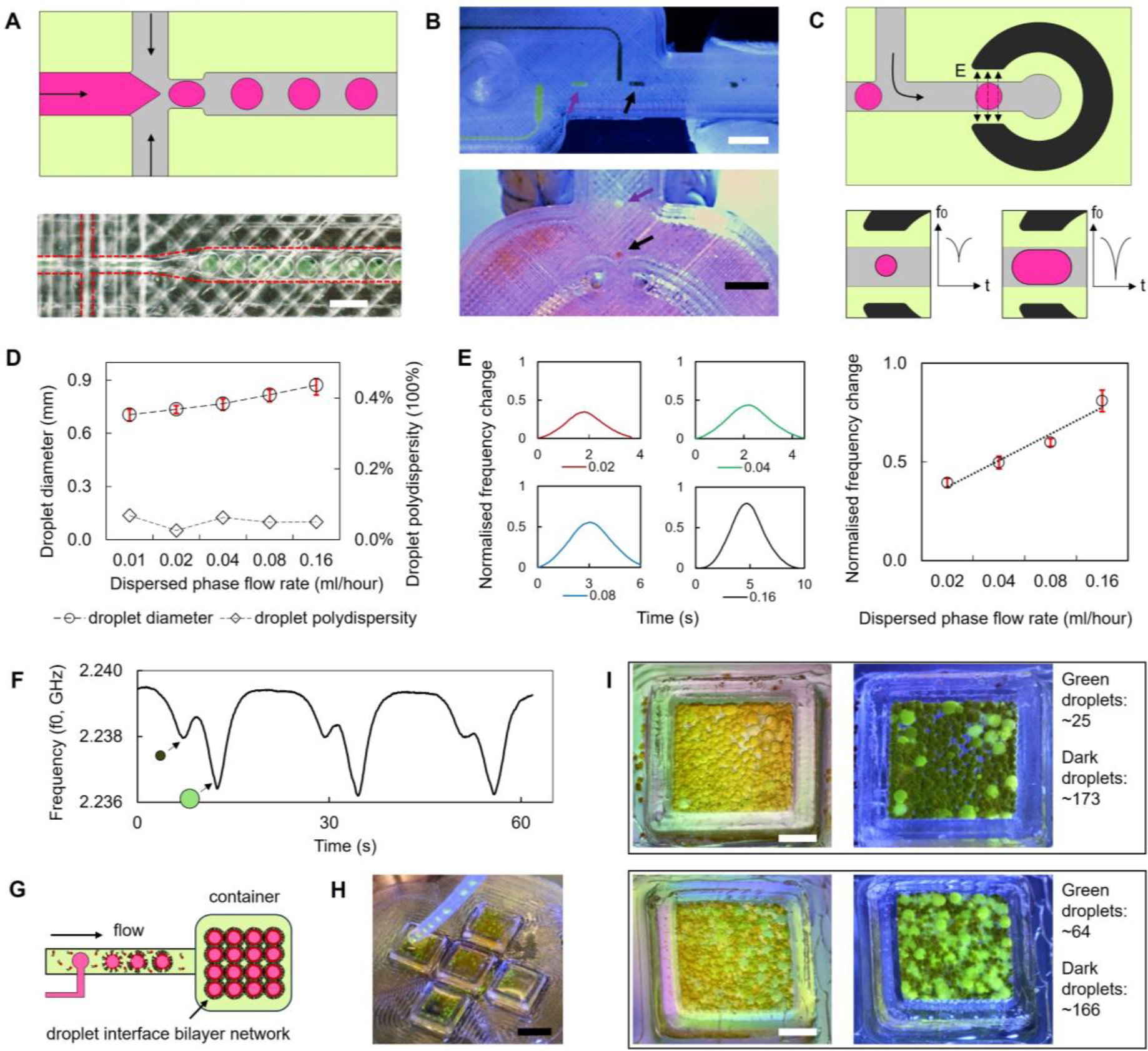
Water-in-oil emulsions and droplet network formation / characterisation, using a 3D-printed MMD. **A**. Droplet generation at a 3D-printed flow-focusing droplet-forming fluidic junctions. Scale bar represents 500 μm. **B**. Droplet generation at 3D-printed T-shaped droplet forming fluidic junctions. Scale bar denotes 5 mm. Green and dark droplets flowed through the gap region of the split-ring microwave resonator. **C**. Larger droplets exhibit higher peaks in the microwave readouts. **D**. Size distribution and polydispersity of droplets under varying flow rates, measured with an optical microscope. The continuous phase flow was maintained at 0.1 ml/hour for all sample groups, with N = 30 for each data point. Error bars indicate the maximum and minimum droplet sizes within the sample groups. Droplet polydispersity was calculated using the formula: polydispersity = ((droplet diameter mean)/ (droplet diameter standard deviation))^2. **E**. Left - Normalised microwave readout frequency for droplets of different sizes corresponding to the input flow rates in Figure 4D. Right – Relationship between droplet diameter and normalised frequencies. Error bars represent frequency variation with N =30 for each data point. **F**. Microwave readout distinguishing small and large droplets, facilitating droplet counting. **G**. Formation of lipid-coated water droplets in a continuous oil phase flow inside a microfluidic duct, leading to DIB network formation when collected in an oil-phase prefilled container. **H**. Aqueous droplets collected in 3D-printed COC petri dishes. Scale bar denotes 1 cm. **I**. Examples of DIB networks with droplets encapsulating different dyes, formed by programmed microfluidic inputs. Approximate droplet counts were derived from the microwave readouts.

### 3D-printed MMD for liquid characterisation

The first type of MMD demonstrated here is designed for analysing liquid samples within a microfluidic chamber. The chamber structure is a cube with an approximate volume of 6 μ*L*, situated at the gap region of the optimised microwave split-ring resonator. Figure 4C depicts the S21 measurement of various samples in these MMDs, including air, an oil mixture, and deionised water (ddH_2_0). All samples showed resonant frequency shifts when alternating between air, water, and oil mixtures in the COC, PP, and PLA MMDs. The oil mixture, comprising 40% silicone oil AR 20 and 60% hexadecane, is an oil phase commonly used in the formation of droplet interface bilayers (DIBs) ^45^. The central resonant frequencies (*f*_0_) for water were approximately 2.1 GHz, 2.25 GHz, and 2.35 GHz, for the PLA, COC and PP MMDs, respectively. Given that 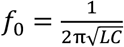, is inversely proportional to the square root of the inductance (L) and capacitance (C), the difference in these frequencies primarily stems from the different material capacitances, as the device geometrical configuration remains constant (capacitance being proportional to material permittivity). Inductance, L is defined by the effective ring length, and capacitance C is affected by the gap distance and cross-sectional area of the liquid metal ring. The distance between the ground plane (i.e., copper film in the MMDs) and the ring is also important.

The effects of liquid metal filling conditions within the 3D-printed channels on MMD sensing performance were assessed. The skin effect in EGaIn at 2 GHz, calculated using 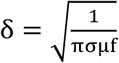, resulted in a skin depth of 6 μm, where σ, μ and *f* are resistivity, relative permittivity, and frequency. To create a thin EGaIn skin within the ring duct, a syringe with needles was employed to expel EGaIn carefully and slowly from the filled ring channel. Fig S3 presents the cross section of the 3D-printed COC channels in various states: empty, fully filled, and with a skin of EGaIn. A residual layer of EGaIn, ranging from 10 μm to 100 μm, adhered to the channel walls, and was likely due to the microstructure created by the FFF 3D-printing process. Microwave sensing performance of the MMD before and after liquid metal removal was tested as shown in Fig S3 table, with air in the gap. The microwave structures resonated under both conditions, indicating effective microwave transmission through the thin EGaIn layer on the COC channel walls. A decrease in quality factor from 101.6 to 48.1 was noted, alongside a slight reduction in central resonance frequency from 2.22 GHz to 2.17 GHz after liquid metal removal and is proposed to be attributed to the heightened current density in the narrow EGaIn layer. Consequently, the air-assisted EGaIn removal had a minimal effect on the resonator sensing performance, except for a marginal decline in measurement resolution due to the decreased Q-factor. Additionally, complete removal of EGaIn from the 3D-printed MMDs was achieved by flushing with a 1 M NaOH solution through the ring structures, which resulted in the disappearance of MMD resonance after complete liquid metal removal.

### 3D-printed MMD for lipid-coated droplets network formation and characterisation

3D-printed microfluidics have been designed to generate complex emulsion droplets^46,47^. The addition of microwave functionality to create MMDs can be used for aqueous droplet formation and processing to non-invasively analyse flowing droplet materials in real-time. As shown in Figure 4A and 4B, flow-focusing droplet-forming fluidic junctions and T-shaped droplet-forming fluidic junctions were printed within the MMD. Downstream from these junctions, an additional fluidic inlet injected an oil phase to regulate the distance between adjacent droplets, ensuring that each droplet passed through the gap region of the split-ring resonator without interference from neighbouring droplets (Figure 4C). The presence of droplet flow can be detected from the microwave readout. Droplet sizes can be analysed in the flow, by volumetric microwave sensing. This is because of the resonant frequency and loss differences between the water and oil materials, with larger aqueous droplets and segments yielding higher peak readings due to their full occupation of the duct.

Droplet sizes were modulated by the input flow rates, as shown in Figure 4D, and the polydispersity of droplets was calculated. Microwave readouts were used to determine the droplet size and polydispersity simultaneously. As shown in figure 4E and 4F, the traces of droplet flow can be observed from the microwave readout, and smaller droplets resulted in shallower troughs on the microwave signatures. The microwave sensing exhibited high sensitivity to the volume variation of droplets, enabling the measurement of the uniformity of microfluidically-formed droplets, as presented in Figure 4G. After calibration of the microwave signatures against optically measured droplet sizes, this technique also characterised droplet populations in water-oil emulsions with diverse components and encapsulants (Figure 4I).Such an approach offers an optical-free method to analyse complex emulsions in situ, which is particularly beneficial for non-transparent microfluidic substrates, like the PLA device shown in Figure 4A, or for liquid samples requiring protection from light exposure. It generates real-time droplet size distribution data for entire droplet populations, eliminating the need for post-sampling droplet collection and physical measurement.

The droplet-forming and sensing functions of the MMDs was employed for the precise construction of lipid-segregated DIB networks. These droplet networks gain functionality through the artificial cellular membranes and reagents encapsulated within each droplet component, for cell-free expression and synthesis ^48^. In our study, DPhPC was pre-dissolved in the oil phase. After forming aqueous droplets in the duct, lipids self-assembled at the water-oil interface, forming a monolayer lipid around the aqueous droplets. Then, the aqueous droplets were collected in a 3D-printed COC petri dish, where the droplets contacted and interconnected to form tissue-like DIB networks, as shown in Figure 4K. The approximate number of droplets was deduced from the microwave readout by calculating the peek values in the traces, allowing the MMD to serve as a non-invasive precision counting tool in this example.

## Discussion

In summary, this study presents a comprehensive study of using fused filament 3D-printing to fabricate microwave-microfluidic devices, incorporating liquid metal injection to form microwave split-ring resonators. We detailed the device designs, printing material selections, and the fabrication and assembly process. The microwave sensing performance of 3D-printed COC MMDs on water-in-oil emulsions was evaluated, yielding results comparable with conventional metal-machined resonators^49^. We demonstrated the application of MMDs for reagent and droplet morphology detection in flow. These findings indicate the efficacy, versatility, and robustness of this fabrication approach for constructing customised microwave-microfluidic devices that can be applied for the non-invasive characterisation of emulsion soft materials.

Most importantly, 3D printing introduces the capability to organise functional fluidic circuits and customised electrical elements on the same substrate, facilitating the complex emulsion formation and processing in one step. This represents a promising engineering tool for soft matter development and construction of bottom-up synthetic cells and tissues. We have show-cased the use of 3D-printed MMDs in determining the compositions of microfluidically formed DIB networks. Future work will integrate this experimental setup with precision droplet deposition methods, such as droplet printing, to analyse in situ droplets distributions and network configurations, enabling studies of networked chemical reactions in macroscale artificial cellular systems.

Although the performance of 3D-printed MMDs is currently constrained by the low resolution of fused filament 3D-printers used here, more accurate and high-resolution 3D-printing platforms, along with more bespoke dielectric materials for microwave and high frequency engineering, are possible^50,51^. The methodology shown here can be significantly improved with these advanced tools to produce customised MMDs and other microelectrode-embedded microfluidic devices featuring complex microstructures. For example, one can conceive the integration of liquid metal electrodes within such duct systems, for electrophysiological investigations, electrocatalytic chemical reactions and the creation of in-duct plasmas.

## Conflicts of interest

There are no conflicts to declare.

## Acknowledgements

DAB, AP, and JL conceived and led the research. KS, JL, PD, CK performed the experiments. JL wrote the manuscript with KS and PD and revised by AP and DAB. This work was partially supported by the European Horizon 2020 project ACDC (Artificial Cells with Distributed Cores) under project award number 824060 and the European Union Horizon Europe EIC 2023 Pathfinder Open programme BioHhOST (bio-hybrid hierarchical organoid-synthetic tissue) under grant agreement No 101130747. The authors also would like to thank Cardiff School of Engineering for the financial support on consumables used in this work. The authors thank Mr. Jacob Fielder, Mr. Ilyaas Rasool, Mr. Daneil Sweeney, Mr. Sameul Bastow, Miss. Chanella Nagra, and Mr. Jake Richardson for their preliminary work on device fabrication.

## Notes

### Competing Interest Statement

The authors have declared no competing interest.

