## Supplementary information for "3D-printing fabrication of microwave-microfluidic device for droplets network formation and characterisation"

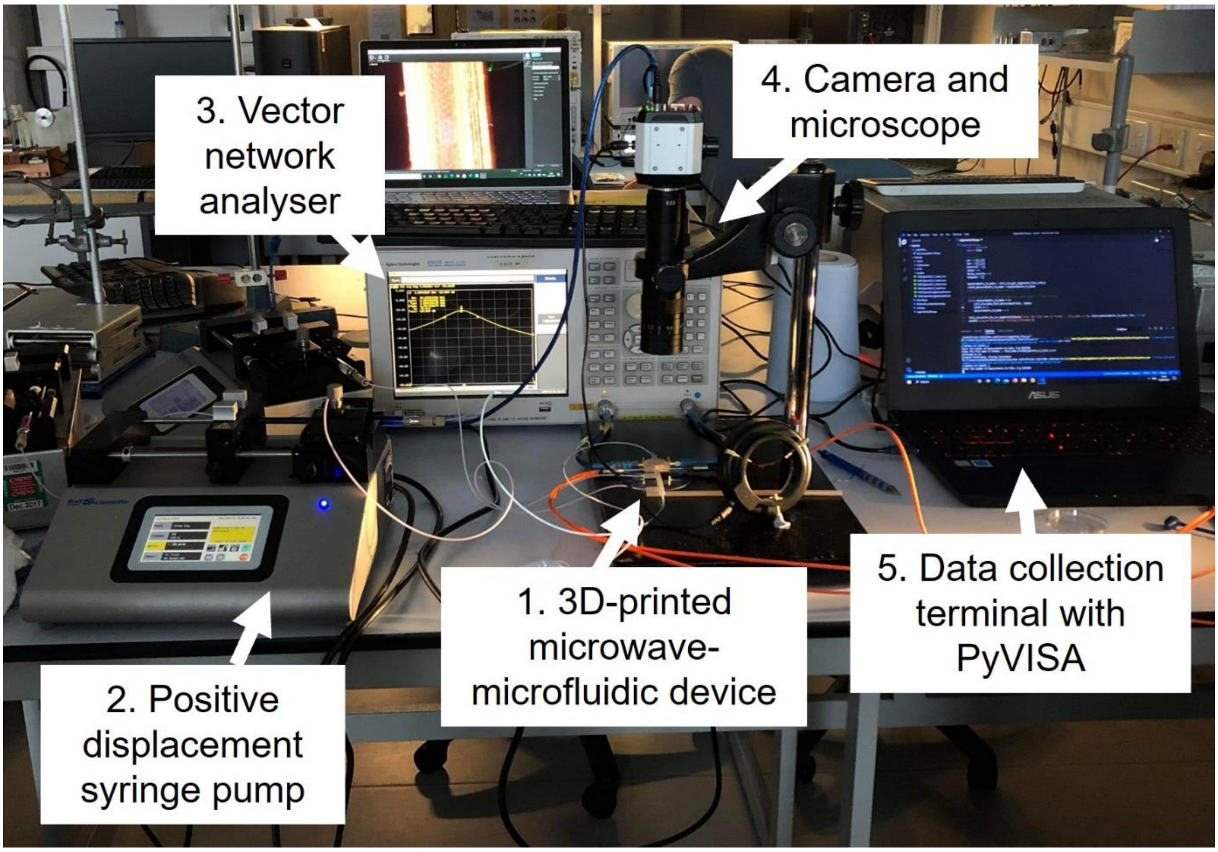


**Fig S1. Experimental setup.** The microwave-microfluidic device was connected to syringe pumps and a vector network analyser for fluid injection and microwave readout, respectively. A CCD camera, attached to an optical microscope, captured the flow of droplets within the microwave-microfluidic device. Microwave readouts were collected using customised code in PyVISA on a Windows laptop.

A

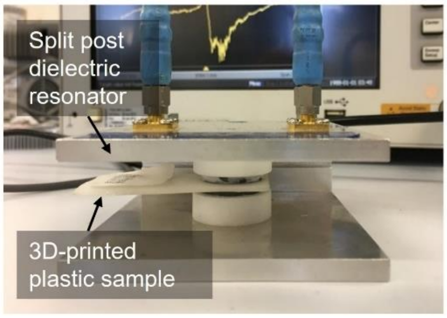


B

| Plastic sample (substrate material) | Printing infill (100%) | $\varepsilon_1$ ±3% | $\varepsilon_2$ ±3% |
|---|---|---|---|
| COC | 100% | 5.35 | 0.010 |
| COC | 50% | 3.63 | 0.020 |
| PLA | 100% | 6.31 | 0.070 |
| Light weight PLA | 100% | 3.95 | 0.042 |
| Nylon | 100% | 7.44 | 0.350 |

**Fig S2. Experimental setup for microwave dielectric characterization of 3D-printed plastic samples. A.** The setup utilises cavity perturbation techniques to measure the voltage transmission coefficient $S_{21}$. A split-post dielectric resonator operating at 2.7 GHz was employed to assess the complex permittivity of the samples. **B.** Table showing the calculated permittivity values of various 3D-printed samples.

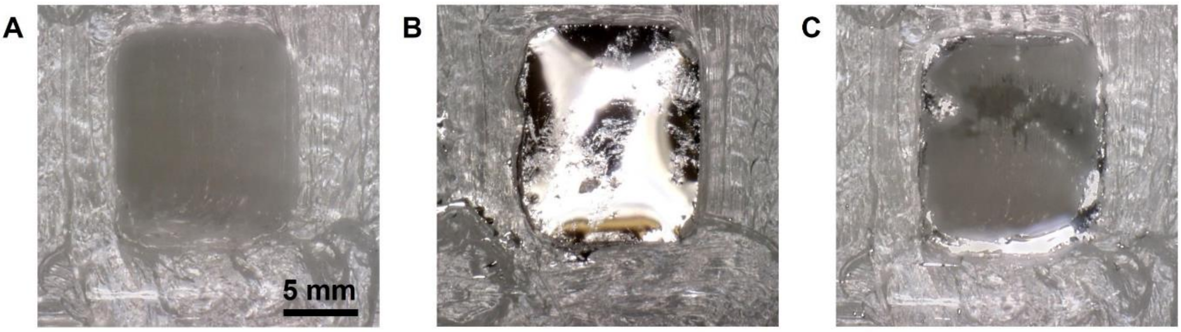


D

| | Bandwidth (MHz) | Centre frequency (GHz) | Quality factor | Transmission loss (dB) |
|---|---|---|---|---|
| Eutectic filled | 21.86 | 2.22 | 101.60 | -9.09 |
| Eutectic removed; thin layer left | 45.05 | 2.17 | 48.07 | -18.41 |

**Fig S3**. **Evaluation of the impact of eutectic filling conditions on MMD performance. A-C.** Sequential cross-sectional views of the ring resonator showing (a) the void channel, (b) channel filled with eutectic materials, and (c) after eutectic removal, with a thin layer of eutectic remaining on the ring channel wall. **D.** Table of microwave parameters for the split-ring microwave resonator before and after eutectic removal. The residual thin layer of eutectic facilitates microwave current transmission, confined to the skin depth, estimated at approximately 6 μmat 2.2 GHz. The microwave resonance is maintained, albeit with a slightly higher resonant frequency and lower Q-factor.
